# scvi-tools: a library for deep probabilistic analysis of single-cell omics data

**DOI:** 10.1101/2021.04.28.441833

**Authors:** Adam Gayoso, Romain Lopez, Galen Xing, Pierre Boyeau, Katherine Wu, Michael Jayasuriya, Edouard Melhman, Maxime Langevin, Yining Liu, Jules Samaran, Gabriel Misrachi, Achille Nazaret, Oscar Clivio, Chenling Xu, Tal Ashuach, Mohammad Lotfollahi, Valentine Svensson, Eduardo da Veiga Beltrame, Carlos Talavera-López, Lior Pachter, Fabian J. Theis, Aaron Streets, Michael I. Jordan, Jeffrey Regier, Nir Yosef

## Abstract

Probabilistic models have provided the underpinnings for state-of-the-art performance in many single-cell omics data analysis tasks, including dimensionality reduction, clustering, differential expression, annotation, removal of unwanted variation, and integration across modalities. Many of the models being deployed are amenable to scalable stochastic inference techniques, and accordingly they are able to process single-cell datasets of realistic and growing sizes. However, the community-wide adoption of probabilistic approaches is hindered by a fractured software ecosystem resulting in an array of packages with distinct, and often complex interfaces. To address this issue, we developed scvi-tools (https://scvi-tools.org), a Python package that implements a variety of leading probabilistic methods. These methods, which cover many fundamental analysis tasks, are accessible through a standardized, easy-to-use interface with direct links to Scanpy, Seurat, and Bioconductor workflows. By standardizing the implementations, we were able to develop and reuse novel functionalities across different models, such as support for complex study designs through nonlinear removal of unwanted variation due to multiple covariates and reference-query integration via scArches. The extensible software building blocks that underlie scvi-tools also enable a developer environment in which new probabilistic models for single cell omics can be efficiently developed, benchmarked, and deployed. We demonstrate this through a code-efficient reimplementation of Stereoscope for deconvolution of spatial transcriptomics profiles. By catering to both the end user and developer audiences, we expect scvi-tools to become an essential software dependency and serve to formulate a community standard for probabilistic modeling of single cell omics.

## 1 Introduction

The field of single-cell omics is rapidly growing, as evidenced by the number of published studies and the number of omics approaches that can reveal distinct aspects of cellular identity [1, 2, 3]. Similar growth has been observed in the number of computational methods designed to analyze single-cell data [4]. These methods overwhelmingly target a core set of computational tasks such as dimensionality reduction (e.g., scVI [5], scLVM [6], CisTopic [7]), cell clustering (e.g., PhenoGraph [8], BISCUIT [9], SIMLR [10]), cell state annotation (e.g., scmap [11], scANVI [12]), removal of unwanted variation (e.g., ZINB-WaVE [13], Scanorama [14], Harmony [15]), differential expression (e.g., DESeq2 [16], edgeR [17]), identification of spatial patterns of gene expression (SpatialDE [18], SPARK [19]), and joint analysis of multi-modal omics data (MOFA+ [20], totalVI [21]).

Many of these methods rely on likelihood-based models to represent variation in the data; we refer to these as *probabilistic models* [22]. Probabilistic models provide principled ways to capture uncertainty in biological systems and are convenient for decomposing the many sources of variation that give rise to omics data [23]. A special class of probabilistic models makes use of neural networks either as part of the generative model, or as a way to amortize the computational cost of inference. These so-called deep generative models (DGMs) have been successfully applied to many analysis tasks for single-cell omics (e.g., scVI [5], totalVI [21], PeakVI [24], SCALE [25], scPhere [26], scGen [27], CellBender [28], Dhaka [29], VASC [30], and scVAE [31]) as well as other areas of computational biology [32].

Despite the appeal of probabilistic models, several obstacles impede their community-wide adoption. The first obstacle, coming from the perspective of the end user, relates to the difficulty of implementing and running such models. As contemporary implementations of probabilistic models leverage modern Python machine learning libraries, users are often required to interact with interfaces and objects that are lower-level in nature than those utilized in popular environments for single-cell data analysis like Bioconductor [33], Seurat [34], or Scanpy [35]. A second obstacle relates to the development of new probabilistic models. From the perspective of developers, there are many necessary routines to implement in support of a probabilistic model, including data handling, tensor computations, training routines that handle device management (e.g., GPU computing), and the underlying optimization, sampling and numerical procedures. While higher-level machine learning packages that automate some of these routines, like PyTorch Lightning [36] or Keras [37], are becoming popular, there still remains overhead to make them work seamlessly with single-cell omics data.

To address these limitations, we built scvi-tools, a Python library for deep probabilistic analysis of single-cell omics data. From the end user’s perspective (Section 2.1), scvi-tools offers standardized access to methods for many single-cell data analysis tasks such as integration of single-cell RNA sequencing (scRNA-seq) data (scVI [5], scArches [38]), annotation of single-cell profiles (CellAssign [39], scANVI [12]), deconvolution of bulk spatial transcriptomics profiles (Stereoscope [40], DestVI [41]), doublet detection (Solo [42]), and multi-modal analysis of CITE-seq data (totalVI [21]). All twelve models currently implemented in scvi-tools interface with Scanpy through the annotated dataset (AnnData [43]) format. Furthermore, scvi-tools has an interface with R, such that all scvi-tools models may be used in Seurat or Bioconductor pipelines.

Here we also demonstrate two new features that add important capabilities for the analysis of large, complex datasets. These two features are accessible for several of the implemented models, and thus, applicable to several types of omics data. The first feature offers the ability to remove unwanted variation due to multiple nuisance factors simultaneously, including both discrete (e.g., batch category) and continuous (e.g., percent mitochondrial reads) factors. We apply this in the context of an scRNA-seq dataset of Drosophila wing development (Section 2.2), which suffered from nuisance variation due to cell cycle, sex, and replicate. The second feature extends several scvi-tools integration methods to iteratively integrate new “query” data into a pre-trained “reference” model via the recently proposed scArches neural network architecture surgery [38]. This feature is particularly useful for incorporating new samples into an analysis without having to reprocess the entire set of samples. We present a case study of applying this approach with totalVI by projecting data from COVID-19 patients into an atlas of immune cells (Section 2.3).

From the perspective of a methods developer, scvi-tools offers a set of building blocks that make it easy to implement new models and modify existing models with minimal code overhead (Section 2.4). These building blocks leverage popular libraries such as AnnData, PyTorch [44], PyTorch Lightning [36], and Pyro [45] and facilitate (deep) probabilistic model design with GPU acceleration. This allows methods developers to focus on developing novel probabilistic models. We demonstrate how these building blocks can be used for efficient development of new models through a reimplementation of Stereoscope, in which a 60% reduction in the number of lines of code was achieved (Section 2.5). This example demonstrates the broad usability and the rich scope of analyses that may be powered by scvi-tools.

## 2 Results

### 2.1 scvi-tools is a unified resource for single-cell omics data analysis

The single-cell omics data analysis pipeline is composed of several steps [46, 47] (Figure 1a). First, data are staged within a data object using packages like Scanpy [35], Seurat [34] or scater [48] and preprocessed with quality control filters. The data are then analyzed through a variety of subsequent steps, which aim to normalize, simplify, infer structure, annotate, and extract new insight (Figure 1b). scvi-tools aims to provide a rich set of methods for these latter steps, while relying on probabilistic models for statistically sound interpretation of data.

**Figure 1:**
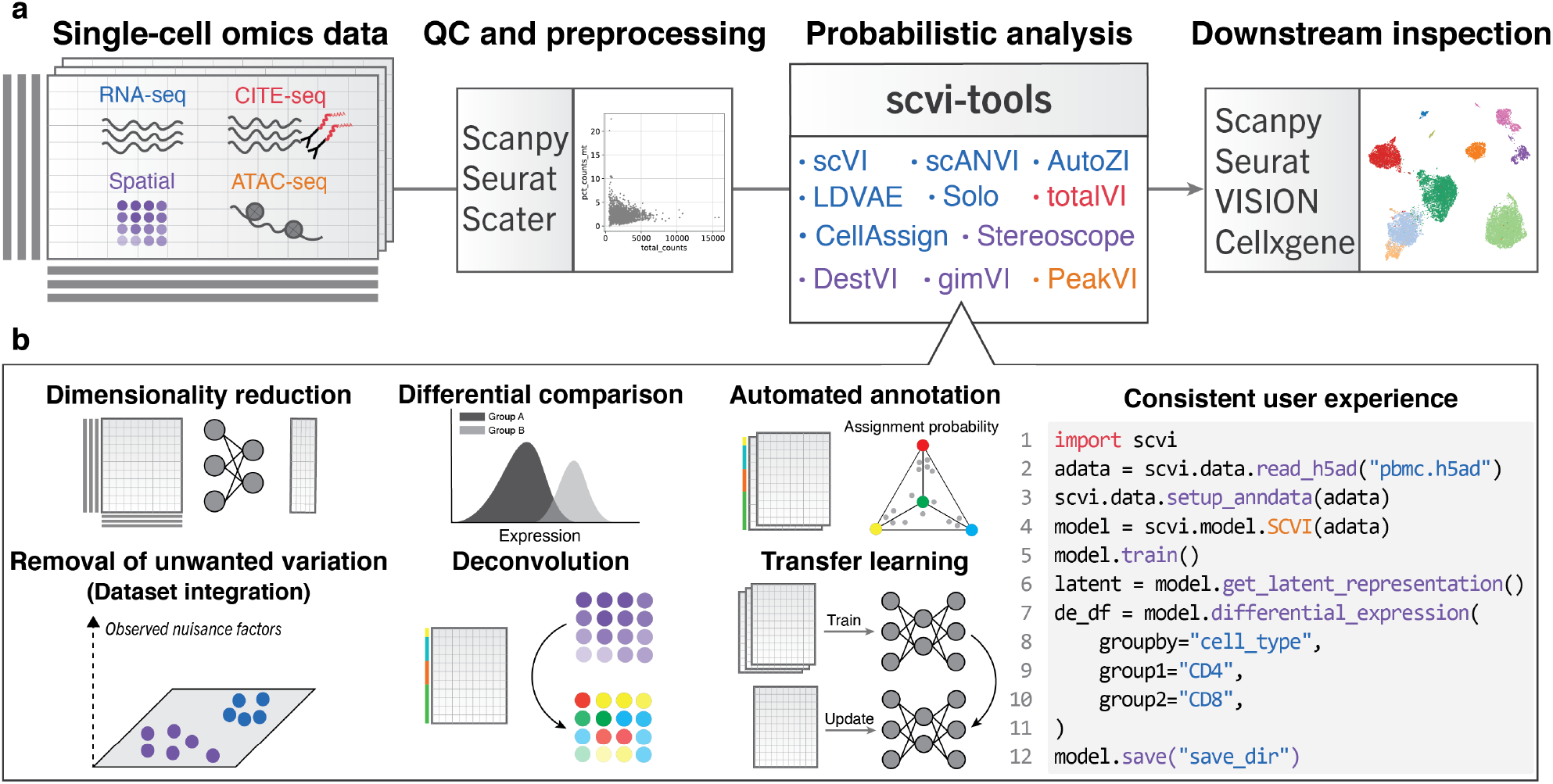
User perspective of scvi-tools. **a**, Overview of single-cell omics analysis pipeline with scvi-tools. Datasets may contain multiple layers of omic information, along with metadata at the cell- and feature-levels. Quality control (QC) and preprocessing are done with popular packages like Scanpy, Seurat, and Scater. Subsequently, datasets can be analyzed with scvi-tools, which contains implementations of probabilistic models that offer a range of capabilities for several omics. Finally, results are further investigated or visualized, typically through the basis of a nearest neighbors graph, and with methods like Scanpy and VISION. **b**, (left) The functionality of models implemented in scvi-tools covers core single-cell analysis tasks. Each model has a simple and consistent user interface. (right) A code snippet applying scVI to a dataset read from a h5ad file, and then performing dimensionality reduction and differential expression.

The models currently implemented in scvi-tools can perform normalization, dimensionality reduction, dataset integration, differential expression (scVI [5, 49], scANVI [12], totalVI [21], PeakVI [24], LDVAE [50]), automated annotation (scANVI, CellAssign [39]), doublet detection (Solo [42]), and deconvolution of spatial transcriptomics profiles (Stereoscope [40], DestVI [41]). These models span multiple modalities including scRNA-seq (scVI, scANVI, CellAssign, Solo), CITE-seq [51] (totalVI), single-cell ATAC-seq (PeakVI), and spatial transcriptomics (Stereoscope, gimVI [52], DestVI [41]) (Supplementary Table 1). Importantly, these models make use of stochastic inference techniques and (optionally) GPU acceleration, such that they readily scale to even the largest datasets.

Each model also comes with a simple and consistent application programming interface (API). They all rely on the popular AnnData format as a way to store and represent the raw data (Figure 1b). Consequently, scvi-tools models are easily integrated with Scanpy workflows. This also enables users to interface with AnnData-based data zoos like Sfaira [53]. Furthermore, the globally consistent API of scvi-tools allows us to maintain a reticulate-based [54] workflow in R, such that scvi-tools models may be used directly in Seurat or Bioconductor workflows. Therefore, after running methods in scvi-tools, results can be visualized and further assessed with a broad range of analysis packages like Scanpy, Seurat, VISION [55], and cellxgene [56].

### 2.2 Nonlinear removal of unwanted variation due to multiple covariates

As the quantity, size, and complexity of single-cell datasets continues to grow, there is a significant need for methods capable of controlling for the effects of unwanted variation [58]. Some factors that contribute to unwanted variation depend directly on the data generating process, such as differences between labs, protocols, technologies, donors, or tissue sites. Normally, such nuisance factors are observable and available as sample-level metadata. Unwanted variation can also come at the level of a single cell and can be calculated directly from the data. It can stem from technical factors like quality as gauged by proxies such as the abundance of mitochondrial RNA or the expression of housekeeping genes, as well as biological factors like cell cycle phase. Nuisance factors can come in either categorical or numerical form. These factors can affect the data in a nonlinear manner [59], and controlling for them is essential for most forms of downstream analysis.

Many methods have been proposed for removing unwanted variation, but most target the subtask of dataset integration, which consists of controlling for one categorical confounding factor at the sample-level, like sample ID [60, 61]. Harmony [15] is, to the best of our knowledge, the only method capable of nonlinearly controlling for multiple categorical covariates simultaneously. While the focus of the integration task is on categorical factors, some pipelines provide an additional layer of normalization, where a given numerical confounder (typically, though not limited to the cell-level) can be regressed out prior to batch correction using a linear model (e.g, the “regress out” function in Scanpy [35] or Combat [62]). Thus, no existing method is capable of performing nonlinear removal of unwanted variation with respect to multiple (cell- or sample-level) categorical and continuous covariates. However, we anticipate an increase in the need for such methods, reflecting the complexity of recent single-cell atlases.

Our previously described models scVI [5], scANVI [12], totalVI [21], and PeakVI [24] all rely on a latent variable models and neural networks to remove unwanted variation from observed covariates, denoted as *s*_*n*_ for a cell *n* (Figure 2a). These models learn a latent representation of each cell that is corrected for this unwanted variation by using a nonlinear neural network decoder that receives the representation and the observed covariates as input, and subsequently injects the covariates into each hidden layer (Figure 2b). While the models could theoretically process multiple categorical or continuous covariates, their previous implementations restricted this capability due to the difficulty of implementing proper data management throughout the training and downstream analysis of a fitted model. scvi-tools allowed us to address this obstacle and support conditioning on arbitrary covariates via its global data registration process and shared neural network building blocks. Now, users simply register their covariates before running a model (Figure 2c). By implementing this feature in the four aforementioned models, scvi-tools can handle complex data collections across a range of modalities and analysis tasks; furthermore, we envision that this feature can be easily propagated to new models (and modalities), all with the same user experience, due to the structured nature of the scvi-tools codebase.

**Figure 2:**
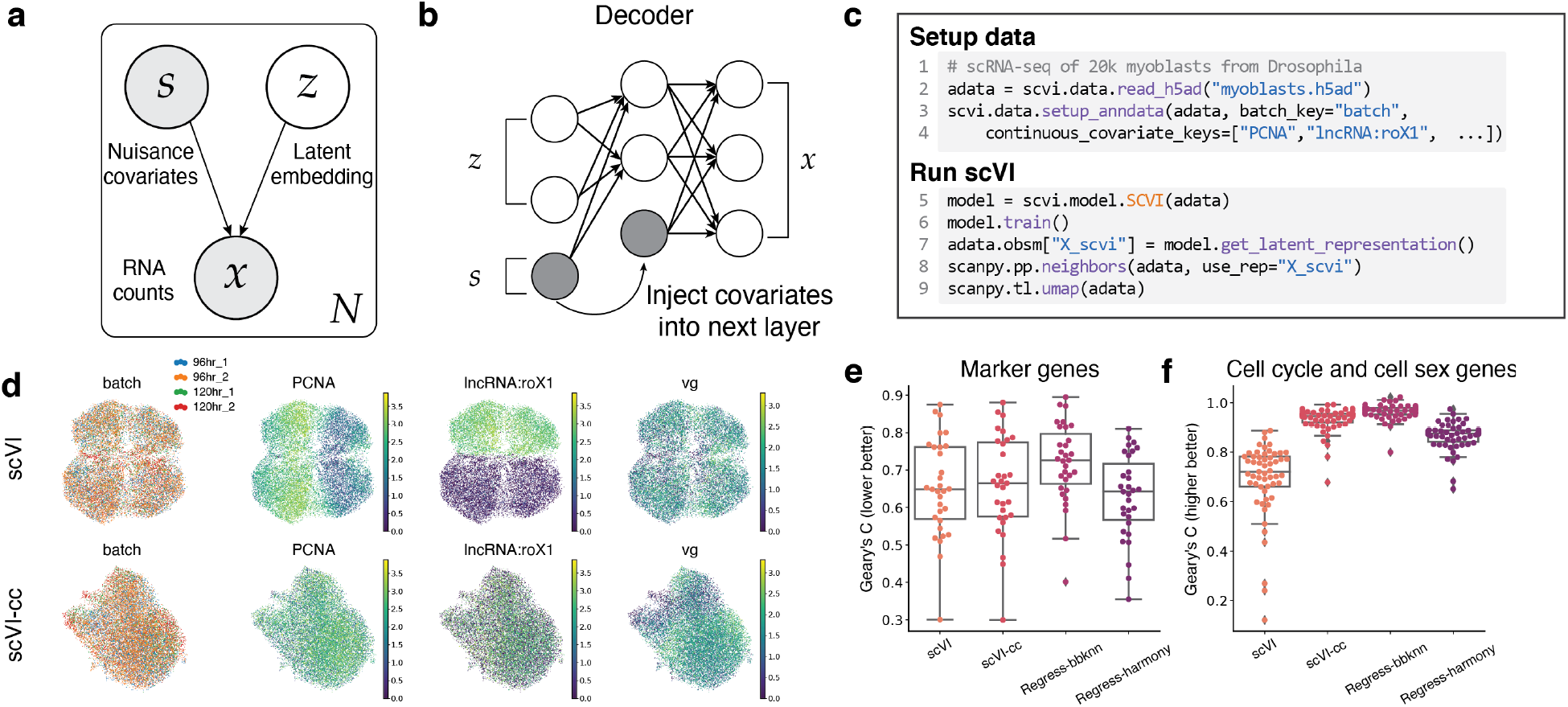
Removal of unwanted variation in the analysis of Drosophila wing disc development. **a**, Graphical model representation of a latent variable models in scvi-tools that conditions on nuisance covariates. **b**, Graphical representation of covariates injected into each layer of decoder neural networks. **c**, Code snippet to register AnnData and train scVI with continuous covariates. The covariates are identified with keys stored in the `AnnData.obs` cell-level data frame. **d**, UMAP [57] embedding of scVI latent space with only batch covariates (scVI) and scVI latent space with batch and continuous covariates (scVI-cc). UMAP plot is colored by batch, *PCNA* (cell cycle gene), *IncRNA:roX1* (cell sex gene), and *vg* (gene marking spatial compartment within the wing disc). **e**, Geary’s C of canonical marker genes of interest per model. **f**, Geary’s C of the cell cycle and cell sex genes conditioned on per model. Box plots were computed on n=31 genes for **(e)** and n=55 genes for **(f)** and indicate the median (center lines), interquartile range (hinges), and whiskers at 1.5× interquartile range. Gene lists can be found in Supplementary Table 2.

To demonstrate this capability, we applied scVI to a dataset of Drosophila wing disc myoblast cells [63] that suffered from strong effects due to cell cycle and sex of the donor organism, both of which were observed via a set of nuisance genes (i.e., numerical covariates; Supplementary Table 2). Both of these confounding factors are observed at the cell-level, as these cells were taken from batches of fly larva and were processed together without sex sorting. The dataset also featured a sample-level confounding factor (batch ID; two batches) and non-nuisance factor (developmental time point; Figure 2d).

When scVI was trained only conditioned on the batch covariate, we observed that it successfully integrated each batch, however the effects from *PCNA*, a cell cycle gene, and *IncRNA:roX1*, a gene that is expressed by males (Figure 2c), still manifested in the latent space. When scVI was additionally conditioned on the expression of the set of nuisance genes (scVI-cc; Methods), we observed that it successfully integrated the data across batch, cell cycle, and sex. Additionally, we found that the desirable biological signals, such as the expression of *vg*, a gene marking a spatial compartment in the wing disc, were preserved (Figure 2d).

We compared these results to the respective Scanpy-based workflow, which consisted of the `scanpy.pp.regress_out` function for linear removal of signal from nuisance genes followed by bbknn [64] (regress-bbknn) or Harmony (regress-harmony) for correction of batch effects (Supplementary Figure 1; Methods). We evaluated these alternative workflows using an autocorrelation measure (Geary’s C [65]) computed with respect to each workflow’s low-dimensional representation and a set of genes, and observed that scVI-cc was able to both retain important biological signal and remove the unwanted variation due to the nuisance genes relative to the vanilla scVI baseline (Figure 2e,f). While we found a tradeoff between retaining biological signal of marker genes and removing nuisance variation across all workflows, these results demonstrate that scVI strikes a balance in removing unwanted variation and retaining wanted variation. The scVI workflow with multiple covariates is also more scalable than the alternatives; we evaluated scalability using subsampled versions of the Heart Cell Atlas dataset [66] that had categorical covariates like donor and continuous covariates like cell-level mitochondrial count percentage (Supplementary Figure 2; Methods).

### 2.3 Transfer learning for reference-query integration with scArches

While dataset integration provides a way to leverage information from many sources, current methods do not scale well to the subtask of reference-query integration in which a “query” dataset is integrated with a large, annotated “reference” dataset. This is an increasingly common scenario, however, that is driven by community efforts for establishing consolidated tissue atlases. These atlases are meant to be used as general references of cell states in a given tissue and may consist of millions of cells [67].

scArches [38] is a recent method that was developed to address this scenario. scArches leverages conditional (variational) autoencoders and transfer learning to decouple the reference-query integration task into two subproblems: First, a reference model is trained on the reference data only; and second, the neural network from the reference model is augmented with nodes that are only influenced by the query data. This new part of the network is subsequently trained with the query data, resulting in a joint model that describes both the train and the reference datasets, while correcting for their technical variation. This procedure dramatically reduces the computational burden of dataset integration. Assuming a pre-trained reference model is available (e.g., representing the “atlas” of cell state for a particular tissue), one only needs to process the (typically much smaller) query data. Another advantage is that the addition of the query data does not change the representation of the reference data in the joint model. Beyond the aforementioned reference-query scenario, this property is also useful in studies where data is accumulated gradually as it eliminates the need for reanalysis when new samples are collected.

We implemented the scArches method in scvi-tools through the addition of one class called `ArchesMixin`. This class contains a generic procedure for adding new nodes to a given reference model, as well as appropriately freezing the nodes corresponding to the reference dataset during training. The `ArchesMixin` class can therefore be inherited by scVI, scANVI, totalVI, and peakVI, without any other custom code. From a user’s perspective, this inheritance adds one function called `load_query_data` to each of these models that is used to load a pre-trained reference model with new query data.

We applied scArches-style reference-query integration with totalVI in order to quickly annotate and then interpret a dataset of immune cells of the blood in donors responding to COVID-19 infection [68] (Figure 3a, Methods). First, we used totalVI to train a reference model of immune cell states, using an annotated CITE-seq dataset of 152,094 peripheral blood mononuclear cells (PBMCs) with over 200 surface proteins [69]. Next, we applied scArches to augment the reference model with an additional CITE-seq dataset of 57,669 PBMCs with 35 proteins from donors with moderate and severe COVID-19, as well as healthy controls [68]. Running totalVI in this setting is made straightforward with the scvi-tools interface and required only a few lines of code (Figure 3a). After query training, we visualized the joint latent representation of the reference and query cells using UMAP (Figure 3b,c). We transferred the annotated cell type labels from the reference cells to the query cells using a random forest classifier operating on the 20 dimensional joint latent space. Notably, since the model does not change for the reference data, the training of the classifier is independent of the query data. Therefore, the classifier itself can also be seen as a part of the pre-trained reference model.

**Figure 3:**
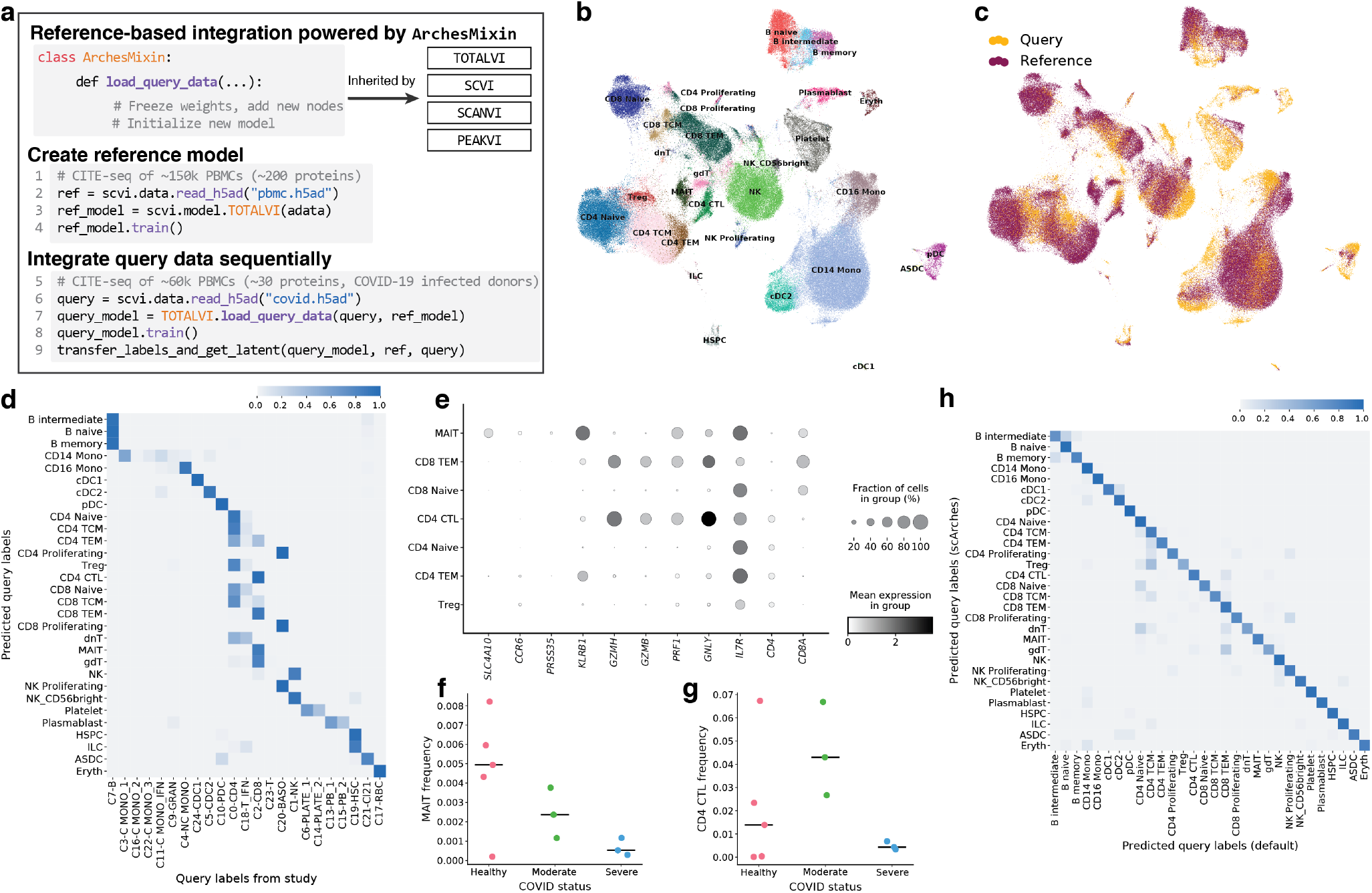
Sequential integration of CITE-seq PBMC samples with totalVI and the scArches method. **a**, Code-based overview of using scArches with the implementation of totalVI in scvi-tools. scArches was implemented globally through the `ArchesMixin` class. First, the reference model is trained on reference data, and then the scArches architectural surgery is performed when `load_query_data` is called on the query data. Finally, the (now) query model is trained with the query data and downstream analysis is performed. **b, c**, UMAP embedding of the totalVI reference and query latent spaces colored by **(b)** the reference labels and predicted query labels and **(c)** the dataset of origin. **d**, Row-normalized confusion matrix of scArches predicted query labels (rows) and study-derived cell annotations (columns). **e**, Dotplot of log library size normalized RNA expression across cell type markers for predicted T cell subsets. **f, g** Frequency of **(f)** MAIT cells and **(g)** CD4 CTLs for each donor in the query dataset across healthy controls and donors with moderate and severe COVID. Horizontal line denotes median. **h**, Row-normalized confusion matrix of scArches predicted query labels (rows) and default totalVI predicted labels (columns).

The predicted labels of the query cells generally agreed with the labels that were provided in the original study and held-out from this analysis (Figure 3d). However, there were some inconsistencies related to T cell subtype classification. The totalVI-scArches approach identified populations of Mucosal associated invariant T (MAIT) cells and CD4 positive cytotoxic T lymphocytes (CD4 CTLs), whereas both populations were mostly annotated as CD8 T cells in the original study (Figure 3d). We found that the predicted MAIT subpopulation had expression of the known markers *SLC4A10, CCR6, KLRB1*, and *PRSS35*, and that they decreased in frequency as a function of COVID severity, which has been previously described [70, 71] (Figure 3e, f). CD4 CTLs have not been as well characterized in terms of their response to COVID, but the predicted CD4 CTLs had relatively high expression of cytotoxic molecules like *PRF1, GZMB, GZMH, GNLY*, and were found to be most prevalent in donors with moderate COVID. This pattern is consistent with evidence suggesting that the presence of CD4 CTLs is associated with better clinical outcomes in other viral infections in humans [72], though more targeted study designs may be necessary to better understand this relationship [73]. Overall, these results suggest that integrating with reference atlases can lead to a more rapid, and potentially more accurate and consistent annotations of cells across studies.

Finally, we compared the totalVI-scArches approach to *default* totalVI, namely training totalVI in one step, using both the reference and query datasets. Using the same random forest procedure, we found that the predictions from scArches and default modes were in high agreement (Figure 3h, Supplementary Figure 3a-e). This is despite the massive speed increase of the scArches approach, which took only 10 minutes to integrate the query dataset, whereas default totalVI took over 80 minutes total due to necessary retraining with the reference data (Supplementary Figure 3h).

### 2.4 scvi-tools accelerates probabilistic model development

scvi-tools has a convenient programming interface for rapid construction and prototyping of novel probabilistic methods, built on top of PyTorch [44] and AnnData [35]. The primary entry point is the `Model` class, which includes all the components needed to fully specify a new probabilistic model. To ensure flexibility, we implemented the `Model` class in a modular manner through four internally used classes (Figure 4a). The `Module` class specifies the probabilistic form of the model (Figure 4b) and contains the elementary calculations that make up the generative model and the inference procedure, including the objective function to optimize during training (e.g., log likelihood or a lower bound thereof). The `TrainingPlan` class defines the procedure for training the model (Supplementary Figure 4a). This class specifies how to manage stochastic gradient descent in terms of optimizer hyperparameters as well as how to update the parameters of a model given random subsamples (i.e., mini-batches) of data. It also provides an interface with PyTorch Lightning’s training procedures [36], which can automatically move the data between different devices such as from CPU to GPU to maximize throughput or perform early stopping. The `AnnDataLoader` class reads data from the AnnData object and automatically structures it for training or for downstream analysis with the trained model (Supplementary Figure 4b). Finally, the `Mixins` are optional classes that implement specific routines for downstream analysis, which can be model-specific or shared among different model classes, such as estimation of differential expression or extraction of latent representations (Supplementary Figure 4c).

**Figure 4:**
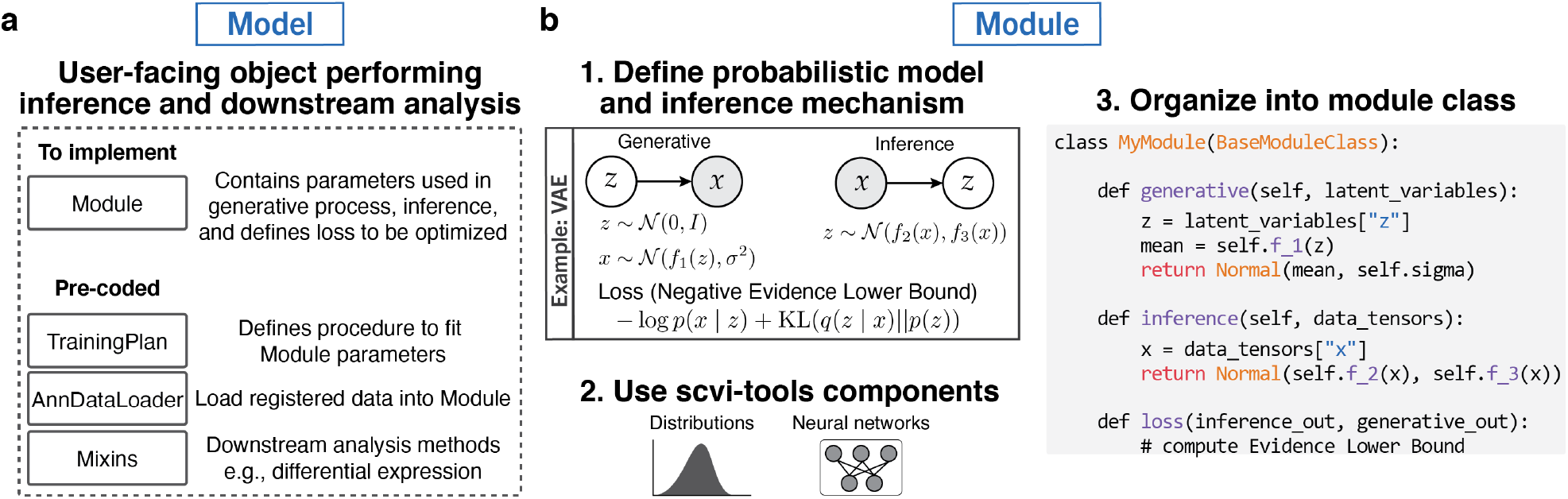
scvi-tools application programming interface for developers. **a**, For every probabilistic method implemented in scvi-tools, users interact with a high-level `Model` object. The Model relies on several lower level components for training a model and analyzing data. The `Module`, which must be implemented systematically, encapsulates the probabilistic specification of the method. The rest of the lower level components rely on pre-coded objects in scvi-tools, such as `AnnDataLoader` for loading data from AnnData objects, `TrainingPlan` for updating the parameters of the module, and `Mixins` classes for downstream analyses. **b**, The creation of a new Module in scvi-tools involves three key steps. First, one mathematically describes the generative model and fully specify the inference procedure. Second, one may choose to from our wide range of pre-coded neural network architectures and distributions, or implement their own in PyTorch object. Finally, those elements are combined together and organized into a class that inherits from the abstract class `BaseModuleClass`. The `generative` method maps latent variables to the data generating distribution. The `inference` method maps input data to the variational distribution (specific to variational inference). The `loss` method specifies the objective function for the training procedure, here the evidence lower bound.

These four components may be reused by many models. The `AnnDataLoader` was written as a generic class and already has support for jointly processing data from multiple modalities, such as transcriptomics and proteomics data in totalVI. The `TrainingPlan` subclasses cover a wide range of scenarios, from optimizing a simple objective function, such as in maximum likelihood estimation (MLE), expectation maximization (EM), or variational inference (VI), to more complex semi-supervised learning procedures as is done in scANVI for handling cells with unobserved annotations. Finally, the available `Mixin` (s), like the `VAEMixin` that offers procedures specific to the variational autoencoder (VAE) (listed in Supplementary Figure 4c), can be inherited by new models and augment their functionalities.

Consequently, methods developers can focus on the `Module` class, which specifies the parameters of the model, the metric of fitness (e.g., data likelihood), and (optionally) a recipe for how latent variables can be sampled given the data. The `Module` class has a generic structure consisting of three functions. First, the `generative` function returns the parameters of the data generating distribution, as a function of latent variables, model parameters, and observed covariates. In the case of a VAE (e.g., scVI, totalVI, etc.), the generative function takes samples of the latent variables as input and returns the underlying data distribution encoded according to the generative (decoder) neural network (Figure 4b). In the case of maximum a posteriori (MAP) inference (as in Stereoscope) or EM (as in CellAssign), the `generative` function maps the model parameters to the data `generating` distribution or returns the components of the expected joint log likelihood, respectively. Second, the `inference` function caters to models that use VI and returns samples from the variational distribution of the latent variables. For a VAE, this calculation is done through an encoder neural network (Figure 4b). Finally, the `loss` function specifies the learning objective given the `inference` and `generative` outputs. For example, the `loss` function can either return the data likelihood (e.g., for MAP), or a lower bound thereof (e.g., for VI or EM).

scvi-tools also contains many pre-coded building blocks that can be used in the development of new `Module` subclasses for new probablistic models. These include popular neural network architectures, as well as distribution classes that are commonly used for single-cell data, like the mean-parameterized negative binomial. scvi-tools also provides an alternative “backend” with Pyro, which has useful properties like automated `loss` computation and (in some cases) inference procedures like automatic differentiation variational inference (ADVI [74]). We provide a more technical exposition of these and other capabilities in the methods section and in tutorials on the scvi-tools website.

### 2.5 Reimplementation of models with scvi-tools

Using the scvi-tools model development interface, we implemented three published methods external to our collaboration: Solo for doublet detection [42], CellAssign for single-cell annotation based on marker genes [39] (Supplementary Note 1), and Stereoscope for deconvolution of spatial transcriptomics profiles [40]. Additionally, we refactored the scGen [27] codebase, a popular method for predicting gene expression perturbations on single cells, to rely on scvi-tools [75]. For all four algorithms, we saw a sharp decrease in number of lines of coded needed. Here we describe the reimplementation of Stereoscope.

Stereoscope is a probabilistic method for deconvolution of spatial transcriptomics profiles, which may represent the average of dozens of cells in each spot [76] (Figure 5a). It is composed of two distinct latent variables models. The first model is trained with an annotated scRNA-seq dataset and learns the gene expression profiles of every annotated cell type. The second model, trained on the spatial data, assumes that the counts in every spot come from a linear combination of the same cell types defined in the scRNA-seq data. The coefficients in this linear combination are normalized and returned as the inferred cell-type proportions at every spot.

**Figure 5:**
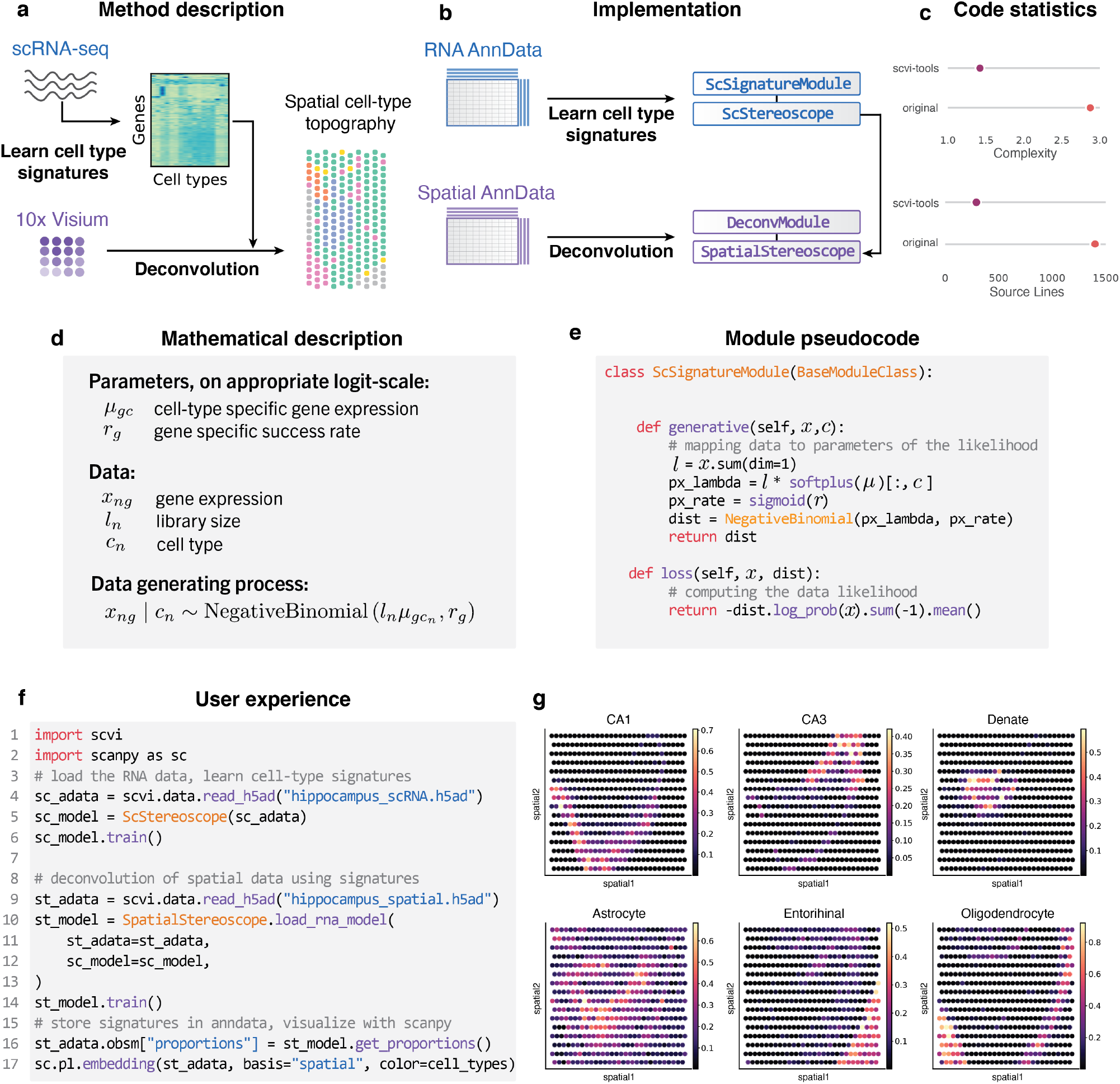
Reimplementation of Stereoscope in scvi-tools. **a**, Overview of the Stereoscope method. Stereoscope takes as input a spatial transcriptomics dataset, as well as single-cell RNA sequencing dataset, and outputs the proportion of cell types in every spot. **b**, Short description of the steps required to reimplement Stereoscope into the codebase. For each of the two models of Stereoscope, we created a module class as well as a model class. **c**, Average cyclomatic code complexity and total number of source code lines for each of scvi-tools implementation and the original implementation. **d, e**, Description of implementation of the `ScSignatureModule`, the module class for the single-cell model of the Stereoscope method. **f**, Example of user code, interaction with Scanpy. **g**, Output example on the hippocampus spatial 10x Visium dataset.

There are several reasons for including Stereoscope in scvi-tools. First, while it is a significant and timely contribution for leveraging spatial transcriptomics, it is difficult to use in practice. Indeed, the reference implementation only provides a command line interface to run the algorithm rather than an API, thus complicating its integration in analysis pipelines. Second, Stereoscope is a linear model that is fit with maximum a posteriori inference. It is therefore conceptually different from many of the other models currently implemented in scvi-tools, most of which are deep generative models trained with amortized variational inference. Consequently, this example illustrates the flexibility of our developer interface. A third reason is the elegance and conciseness of this model, which made it a good case study for demonstrating an implementation with our PyTorch backend.

Using the scvi-tools developer interface provided both a conceptual and practical simplification of the reimplementation, focusing most of the effort on the formulation of the actual probabilistic model. Specifically, our implementation consisted of two module classes and two model classes (one pair of classes per latent variable model; Figure 5b). It was not necessary to write any code for data loading or training, as these functionalities are inherited through the scvi-tools base classes. Consequently, we observed a marked reduction both in the code complexity (average cyclomatic complexity [77]) and the number of lines of code (Methods) compared to the original codebase (Figure 5c).

To further illustrate the simplicity of implementing new models in scvi-tools, we elaborate on the development of the `ScSignatureModule` class (the module class for the scRNA-seq data latent variable model). In the model, for every cell *n*, the observed data includes its cell type *c*_*n*_, its library size *l*_*n*_ and for every gene *g*, the gene expression *x*_*ng*_. Let *G* be the number of observed genes and let *C* be the number of annotated cell types. The parameters of the model, which we want to infer, specify the distribution of each gene *g* in every cell type *c*. This distribution is assumed to be a negative binomial, parameterized by λ_*ng*_ and *r*_*g*_, where (*r*_*g*_*λ*_*ng*_)*/*(1 − *r*_*g*_) is the expectation and where *r*_*g*_ is a gene-specific parameter determining its mean-variance relationship. Parameter 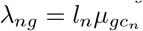 of the negative binomial depends on the type assigned to the cell (*c*_*n*_), its overall number of detected molecules (*l*_*n*_). Figure 5d specifies the model more concisely.

The parameters of the data generative procedure are calculated in `ScSignatureModule` (Figure 5e). While the code follows the model closely, care must be given to the constraints on the parameters. For example, *µ* must be positive, so we use a softplus transformation. Similarly, *r* must be in the range [0, 1], which we enforce with a sigmoid transformation. The loss function returns the likelihood of the observed data, using a negative binomial distribution and evaluated at the model parameters. In contrast to VAEs (Figure 4b), there is no need to provide an implementation of the inference method because there are no latent variables. Our implementation therefore consists only of the generative function and the loss.

We applied the method to the 10x Visium spatial transcriptomics data of an adult mouse brain [78] and a single-cell RNA sequencing dataset of the mouse hippocampus [79] (Methods). A schematic of the user experience in Figure 5f. Applying Stereoscope and visualizing the results with Scanpy takes less than 20 lines of code including the import statements and the call to the Scanpy library. In contrast to the original software, which only accommodated a command line interface, our reimplementation can be used from Jupyter notebooks and with AnnData objects. The result of our Stereoscope implementation, which appears in Figure 5g, performs nearly the same as the original implementation (Supplementary Figure 5) in terms of the average Spearman correlation of cell-type proportion across all individual cell types.

## 3 Discussion

We have developed scvi-tools, a resource for the probabilistic analysis of single-cell omics data with standardized access to more than ten models that can collectively perform a multitude of tasks across several omics. We also equipped several models with new features to aid in the analysis of large, complex data collections via the implementations of removal of unwanted variation due to arbitrary covariates and reference-query integration with scArches. To simplify the usage of scvi-tools, we made the functionalities of models concentrated in scope and ensured seamless connection with the single-cell software ecosystem at large. Thus, preprocessing and downstream analysis (e.g., clustering of a latent space) can be performed with widely-used packages like Scanpy and Seurat. On the scvi-tools documentation website, we feature the API reference of each model as well as tutorials describing the functionality of each model and its interaction with other single-cell tools. We also made these tutorials available via Google Colab, which provides a free compute environment and GPU and can even support large-scale analyses.

At its core, scvi-tools is based on reusable software building blocks for constructing single-cell-focused probabilistic models. With these building blocks, we reimplemented our own previously described models and external models, which together span several inference techniques, like MAP inference, EM, and many stochastic variational inference algorithms [80] such as amortized variational inference [81]. Further, standardized implementations of these models allowed us to add features to many models simultaneously, like in the case of scArches. We expect that this type of method development will grow in popularity following future advancements in microfluidics and molecular biology that will produce a new set of computational challenges.

scvi-tools models contain modules that structure the implementation of a generative process and inference procedure. While we focused on the developing modules directly with PyTorch, we can alternatively develop a module with Pyro. From a developer’s perspective, it may be unclear when one should use the PyTorch backend or the Pyro backend for module development. Using Pyro comes with many benefits, such as automated ELBO computation and automatic differentiation variational inference [74]. We envision that Pyro will be useful for hierarchical Bayesian models with a large collection of latent variables, such as cell2location [82]. Pyro will be also appropriate in cases where a black-box inference procedure is known to work well. However, for more complex cases with inference schemes that deviate from standard Bayesian recipes (e.g., warmup [83], alternate variational bounds [84, 85], adversarial inference [86]), we recommend using the PyTorch backend.

In the development of scvi-tools, we aimed to bridge the gap that exists between the single-cell software ecosystems and the contemporary machine learning frameworks for constructing and deploying this class of models. Thus, developers can now expect to build models using popular machine learning libraries that are immediately accessible to end users in the single-cell community. On our documentation website, we provide a series of tutorials on building a model with scvi-tools, walking through the steps of data management, module construction, and model development. We also built a template repository on GitHub that enables developers to quickly create a Python package that uses unit testing, automated documentation, and popular code styling libraries. This repository demonstrates how the scvi-tools building blocks, through a simplified implementation of scVI, can be used externally. We anticipate that most models built with scvi-tools will be deployed in this way as independent packages.

scvi-tools remains under active development. Thus, end users can expect that scvi-tools will continually evolve along with the field, adding support for new models, new workflows, and new features. Looking forward, we anticipate that these resources will serve the single-cell community by facilitating the prototyping of new models, creating a standard for the deployment of probabilistic analysis software, and enhancing the scientific discovery pipeline.

## Acknowledgements

We acknowledge members of the Streets and Yosef laboratories for general feedback. We thank all the GitHub users who contributed code to scvi-tools over the years. We thank Nicholas Everetts for help with the analysis of the Drosophila data. We thank David Kelley and Nick Bernstein for help with implementing Solo. We thank Marco Wagenstetter and Sergei Rybakov for help with the transition of the scGen package to use scvi-tools as well as feedback on the scArches implementation. We thank Hector Roux de Bézieux for insightful discussions about the R ecosystem. We thank Kieran Campbell and Allen Zhang for clarifying aspects of the original CellAssign implementation. We thank the Pyro team, including Eli Bingham, Martin Jankowiak, and Fritz Obermeyer, for help with integrating Pyro in scvi-tools. Research reported in this manuscript was supported by the NIGMS of the National Institutes of Health under award number R35GM124916 and by the Chan-Zuckerberg Foundation Network under grant number 2019-02452. A.G. is supported by NIH Training Grant 5T32HG000047-19. A.S. and N.Y. are Chan Zuckerberg Biohub investigators.

## Author contributions

A.G., R.L, and G.X. contributed equally. A.G. designed the scvi-tools application programming interface with input from G.X. and R.L. G.X. and A.G. lead development of scvi-tools with input from R.L. G.X. reimplemented scVI, totalVI, AutoZI, and scANVI with input from A.G. R.L. implemented Stereoscope with input from A.G. Data analysis in this manuscript was led by A.G., R.L., and G.X with input from N.Y. A.G, R.L, P.B, E.M, M.L, Y.L, J.S, G.M, A.N, O.C. worked on the initial version of the codebase (scvi package), with input from M.I.J, J.R and N.Y. R.L, E.M and C.X contributed the scANVI model, with input from J.R and N.Y. A.G implemented totalVI with input from A.S and N.Y. T.A. implemented peakVI with input from A.G. A.G implemented scArches with input from M.L, F.T and N.Y. V.S. made several contributions to the codebase, including the LDVAE model. P.B. contributed the differential expression programming interface. E.B and C.T.L provided tutorials on differential expression and deconvolution of spatial transcriptomics, with input from L.P. K.W implemented CellAssign in the codebase with input from A.G.. M.J. made general code contributions and helped maintain scvi-tools. N.Y. supervised all research. A.G, R.L, G.X, J.R and N.Y wrote the manuscript.

## Ethics declaration

O.C. is supported by the EPSRC Centre for Doctoral Training in Modern Statistics and Statistical Machine Learning (EP/S023151/1) and Novo Nordisk. V.S. is a full-time employee of Serqet Therapuetics and has ownership interest in Serqet Therapeutics. F.J.T. reports receiving consulting fees from Roche Diagnostics GmbH and Cellarity Inc., and ownership interest in Cellarity, Inc.

## 4 Methods

scvi-tools is a Python package available via the Python Package Index (PyPI) and Bioconda. Further details on scvi-tools, including model vignettes and a tutorial series on developing models, are available at https://scvi-tools.org.

### Analysis with multiple covariates

#### Drosophila

The dataset of Drosophila myoblasts was downloaded from Gene Expression Omnibus (Accession ID GSE155543). The list of cell cycle, cell sex, and marker genes were provided by the authors (Supplementary Table 2). scVI was trained on the raw count data with a single batch covariate for timepoint and replicate. For scVI-cc, the model was conditioned on each nuisance gene’s log normalized expression (first each cells counts were normalized, then scaled to the median of total counts for cells before normalization, before calculating the log on the scaled normalized counts) as well as a categorical batch covariate for timepoint and replicate. Furthermore, each nuisance gene that was conditioned on was also removed from the input count matrix. Each scVI model was trained for 400 epochs. For regress-bbknn, the `regress_out` scanpy function was used for the continuous covariates, while the `bbknn` scanpy function was used for the categorical batch covariates. Similar to scVI-cc, the nuisance gene expression was removed from the input count matrix. For regress-harmony, the `regress_out` function was used for the continuous covariates, while the `harmony_integrate` scanpy function was used for the categorical batch covariates. The nuisance genes were removed from the input count matrix. To compute Geary’s C we used the `gearys_c` scanpy function.

#### Runtime

Runtime analysis was run on a desktop with an Intel Core i9-10900K 3.7 GHz processor, 2x Corsair Vengeance LPX 64GB ram, and an NVIDIA RTX 3090 GPU. Runtime was performed with the Heart Cell Atlas dataset downloaded from https://www.heartcellatlas.org/. We then subsampled and created datasets of 5,000, 10,000, 20,000, 40,000, 80,000, 160,000, 320,000, and 486,134 cells and selected the top 4,000 genes via `highly_variable_genes` per dataset, with parameter `flavor=“seurat_v3”`. For each of the following methods, we used the cell_source and donor fields of the dataset as categorical covariates. For continuous covariates, we generated 8 random covariates by sampling from a standard normal distribution in addition to treating the percent_mito and percent_ribo fields as additional continuous covariates, for a total of 10 continuous covariates. For scVI runtime, we tracked the runtime of running the `train` function with the following parameters:

~~~
early_stopping=True, early_stopping_patience=45, max_epochs=10000,
batch_size=1024, limit_train_batches=20,
train_size=0.9 if n_cells < 200000 and train_size=1-(20000/n_cells) otherwise
~~~

This choice of the size parameter for the training set ensures that the validation set always has size of less than 20,000 cells for the whole runtime experiment. For the regress-bbknn baseline, we tracked the runtime of correcting continuous covariates with `regress_out,` running pca, and correcting for categorical covariates with `bbknn` (all from the scanpy package). For the regress-harmony baseline, we tracked the runtime of `regress_out` to correct for continuous covariates, running `pca,` and correcting for categorical covariates with `harmony_integrate` (all from the scanpy package). For the regress baseline, we only tracked the runtime of the `regress_out` function.

### Transfer learning with scArches

The reference dataset of Human PBMCs corresponding to the CITE-seq dataset described in [69] was downloaded from Gene Expression Omnibus (Accession ID GSE164378). Associated metadata, like cell-type annotations, were retrieved from https://atlas.fredhutch.org/nygc/multimodal-pbmc/. The query dataset of Human PBMCs corresponding to the CITE-seq dataset described in ref. [68] was provided by the authors, with full metadata including study-derived annotations and other study design information. The reference data were additionally filtered to include cells that meet the following criteria: (1) greater than 150 proteins detected, (2) percent mitochondrial counts less than 12%, (3) not doublets (i.e., reference cells annotated as doublets removed), (4) natural log protein library size, defined as total protein counts, between 7.6 and 10.3. Furthermore, protein features corresponding to isotype controls were removed (for reference and query), and protein features targeting the same protein (i.e., antibody clones) were summed together (applies to reference only). The query data were additionally filtered for doublets using Scrublet [87] with default parameters. Highly variable genes (4000) were selected using only the reference datasets and with the method in Scanpy `(flavor=“seurat_v3”)`. Finally, the protein expression for a random set of five out of the 24 batch categories (representing time and donor) in the reference dataset were masked from totalVI, in order to help the model generalize to query data with mismatched proteins.

In the case of scArches, totalVI was trained on the reference data for 250 epochs, using two hidden layers and layer normalization as described in the totalVI scArches tutorial https://docs.scvi-tools.org/en/stable/user_guide/notebooks/scarches_scvi_tools.html. After applying the scArches architectural surgery, the model was updated with the query data for 150 additional epochs. The latent representation for the reference and query datasets was obtained from the model after it was updated with the query data; however, we note that the latent representation of the reference data does not change after query traning in the case of scArches. In the case of default totalVI, we kept all hyperparameters and data preprocessing the same, except that we trained totalVI only once, on the concatenated reference and query datasets for 250 epochs.

For both scArches and default totalVI, we transferred the reference cell-type annotations using a random forest classifier implemented in scikit-learn [88]. In particular, we trained a random forest classifier on the latent representations of the reference cells (default parameters except `class_weight=“balanced_subsample”)`. The query cell annotations were then obtained by passing the query latent representations to the classifier. In all cases, UMAP embeddings were computed using Scanpy [89] with `metric=“cosine”` and `min_dist=0.3`. The frequency of predicted query MAIT cells and CD4 CTLs were computed on a per-donor basis and were defined as the frequency given the total number of observed cells per donor. All analyses were run on a computer with one NVIDIA GeForce RTX 3090 GPU.

### Cell-type deconvolution with Stereoscope

For the single-cell data, we used the dataset from Saunders et al. [79], as pre-processed by Cable et al [90]. We filtered genes out with a minimum count of 10. Then, we selected 2,000 highly variable genes using the corresponding scanpy function. For the spatial transcriptomics data, we used the V1 Adult Mouse Brain dataset [78] and filtered spots so as to focus on the hippocampus (as in Cable et al. Supplementary Figure 7). We then filtered genes in the spatial transcriptomics data by taking the intersection with the highly variable genes in the single-cell data. We then ran the single-cell model for 100 epochs and ran the spatial model for 5,000 epochs. We ran the original Stereoscope code from the command line, on the same dataset and with the same parameters. We used the Radon package for the calculations of average cyclomatic complexity [77] and source lines of code.

## Data availability

A collection of processed data discussed in this manuscript have been deposited on figshare (https://doi.org/10.6084/m9.figshare.14374574.v1).

## Code availability

The code to reproduce the experiments of this manuscript is available at https://github.com/YosefLab/scvi-tools-reproducibility. The `scvi-tools` package can be found on GitHub at https://github.com/YosefLab/scvi-tools, and is also deposited on Zenodo https://doi.org/10.5281/zenodo.4341715.

## Supplementary Figures

**Supplementary Figure 1:**
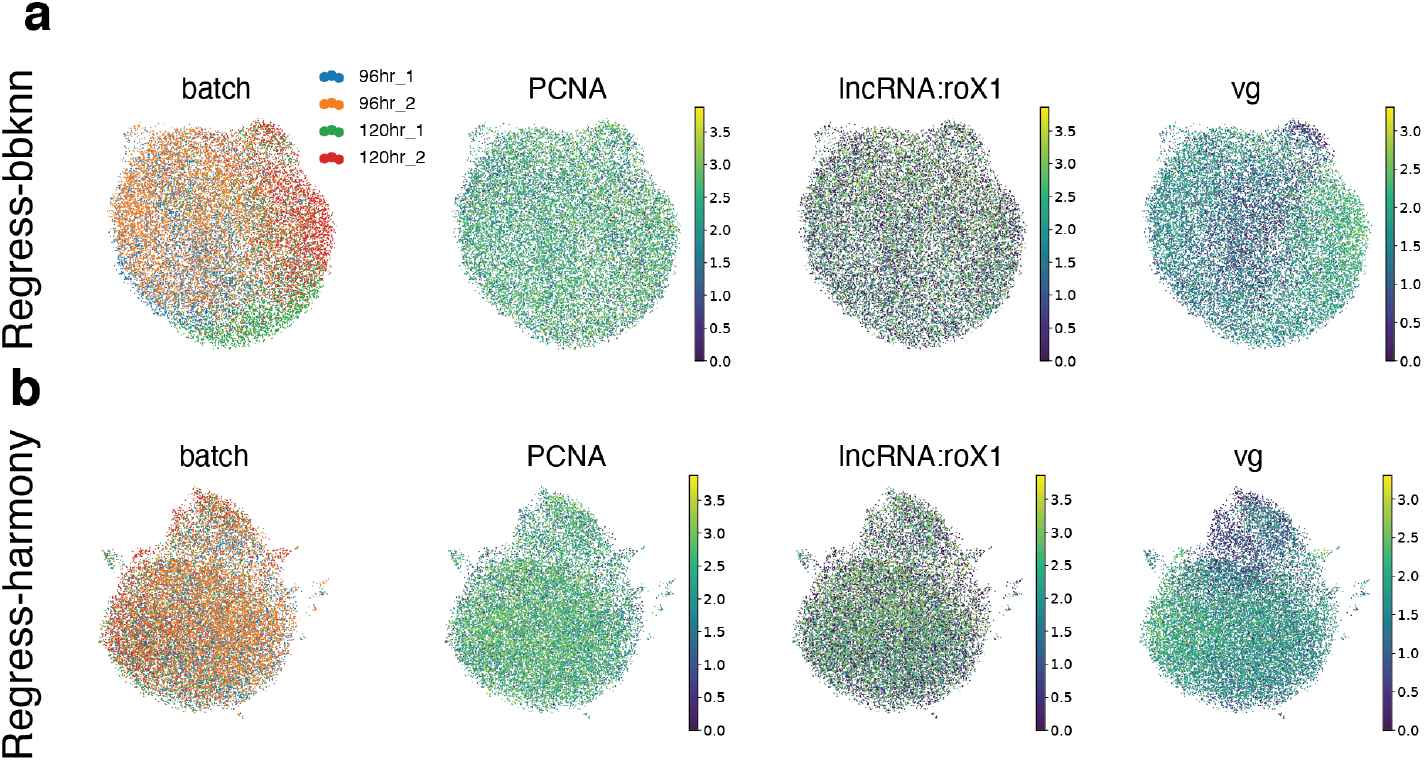
Multiple covariates. **a, b**, Low-dimensional embeddings derived from `scanpy.pp.regress_out` function followed by **(a)** Scanpy’s external bbknn and **(b)** harmony methods (Regress-bbknn, Regress-harmony; Methods). UMAP plot is colored by batch, *PCNA* (cell cycle gene), *IncRNA:roX1* (cell sex gene), and *vg* (gene marking spatial compartment within the wing disc).

**Supplementary Figure 2:**
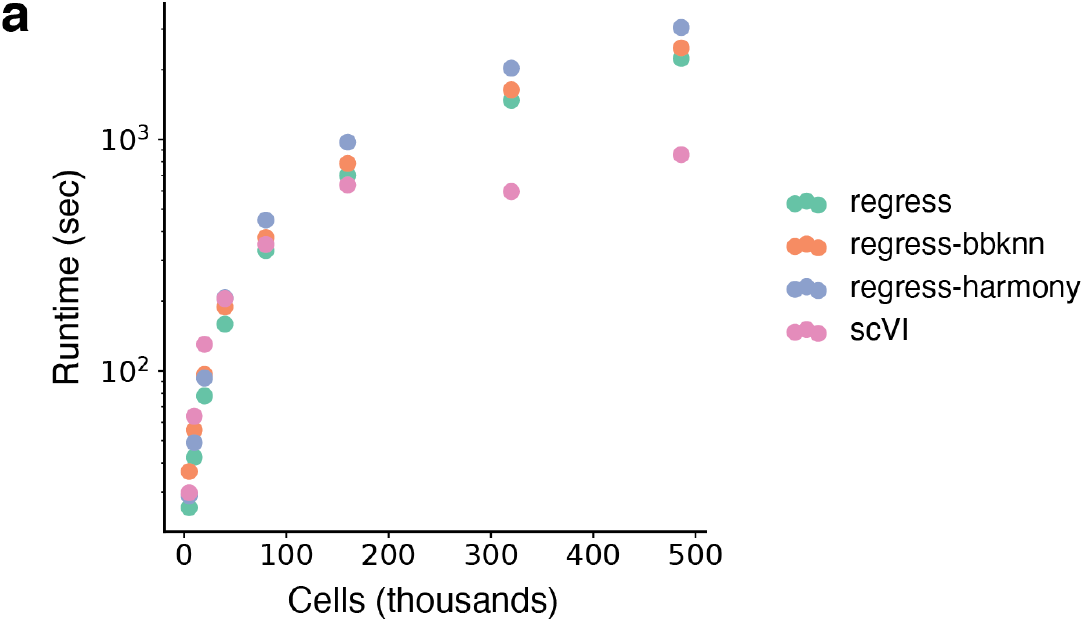
Multiple covariates runtime on Heart Cell Atlas. **a**, Runtime of workflows to correct multiple continuous and categorical covariates on subsampled versions of the Heart Cell Atlas dataset.

**Supplementary Figure 3:**
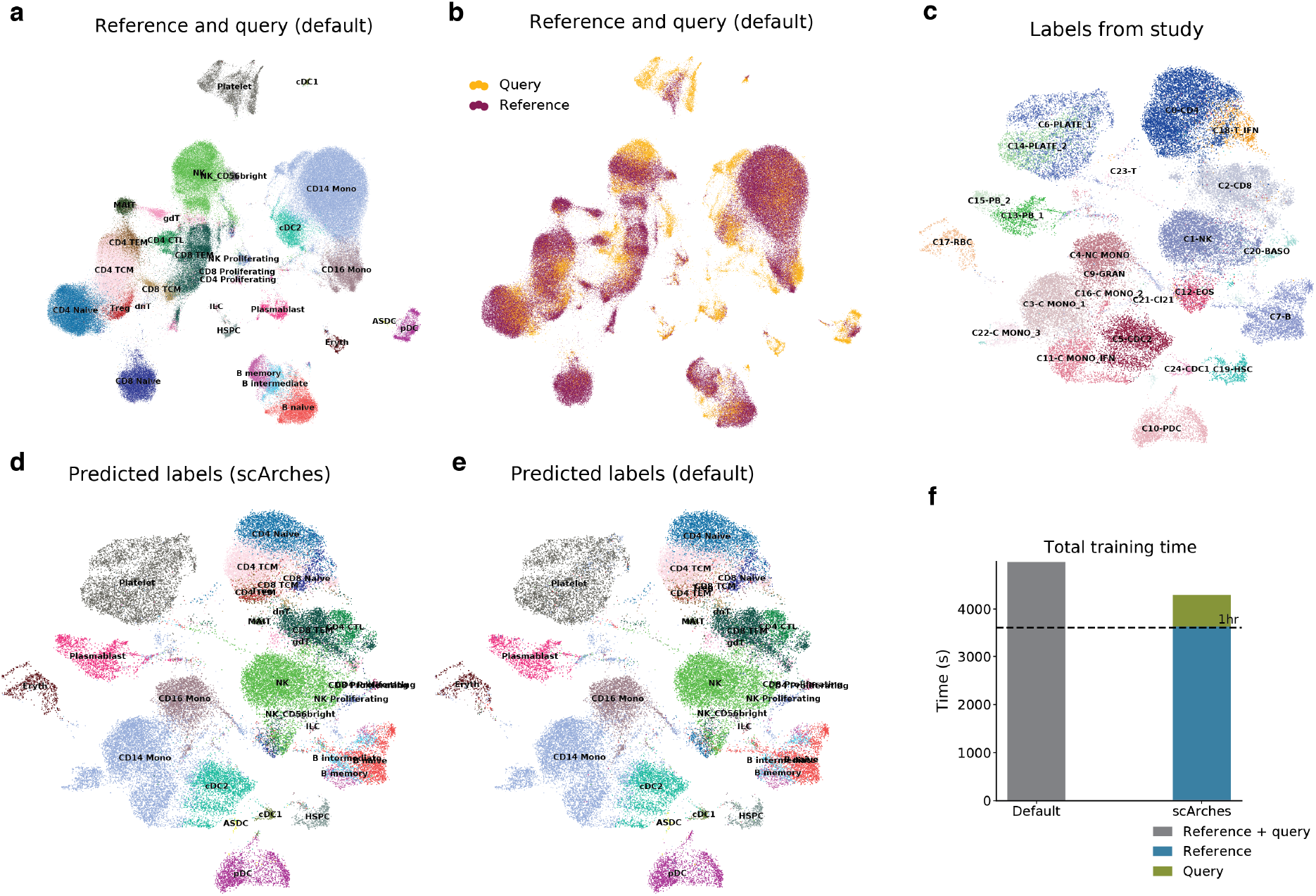
Sequential integration of PBMC samples with totalVI and the scArches method. **a, b**, UMAP embedding of the totalVI reference and query latent spaces from default totalVI model run colored by **(a)** the reference labels and predicted query labels and **(b)** the dataset of origin. **c-e**, UMAP embedding of the totalVI scArches query latent space from colored by **(c)** study-derived labels, **(d)** predicted labels from totalVI scArches, and **(e)** predicted labels from totalVI default. **f**, Runtime of totalVI default versus totalVI scArches.

**Supplementary Figure 4:**
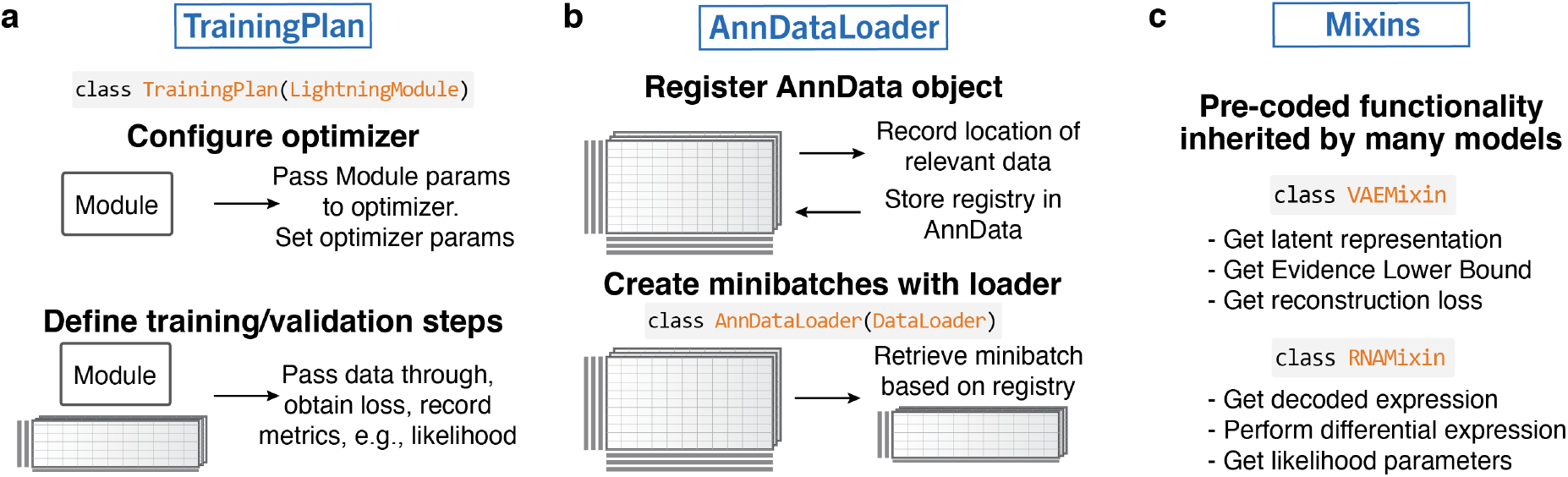
Black box components of scvi-tools’ application programming interface for developers. **a**, The training plan configures several aspects relevant for model training, like the optimizers, and the actual training optimization step in which data gets passed through a module. **b**, The `AnnDataLoader` is used to load minibatches of data directly from an AnnData object into a module. **c**, `Mixins` are Python classes with pre-coded functionality that can be inherited into many models, introducing isolated feature sets, like getting metrics relevant to VAEs (VAEMixin).

**Supplementary Figure 5:**
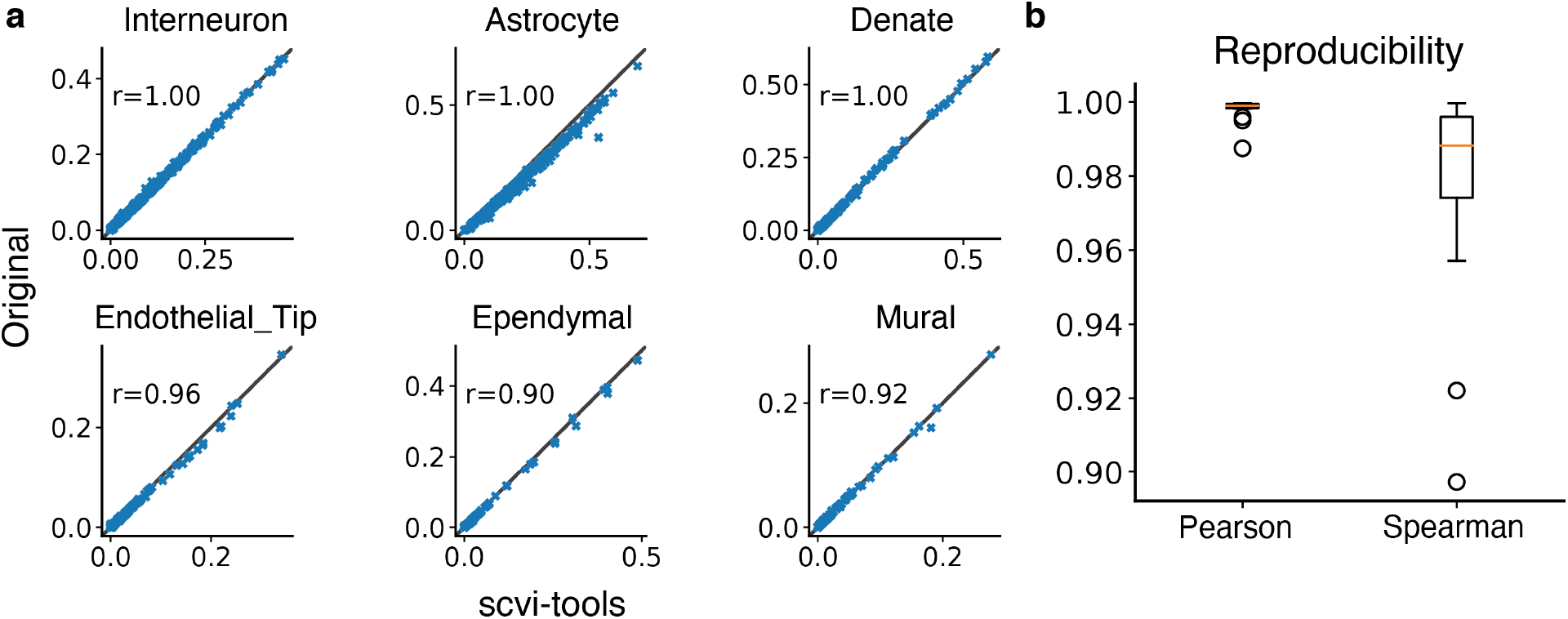
Reproducibility of Stereoscope implementation in scvi-tools. **a**, Scatter plot for six cell types of the hippocampus dataset. Each point in the scatter plot represents one spot. The *x* axis is the proportion inferred by the original Stereoscope software and the *y* axis is the proportion inferred by our implementation. We report the Pearson coefficient for each cell type. The top three cell types are the most reproducible and the bottom three cell types are the least reproducible. **b**, Box plot of correlations of proportions between the two implementations. Each dot in the box plot is a cell type. Golden line in the box plot is the median.

**Supplementary Figure 6:**
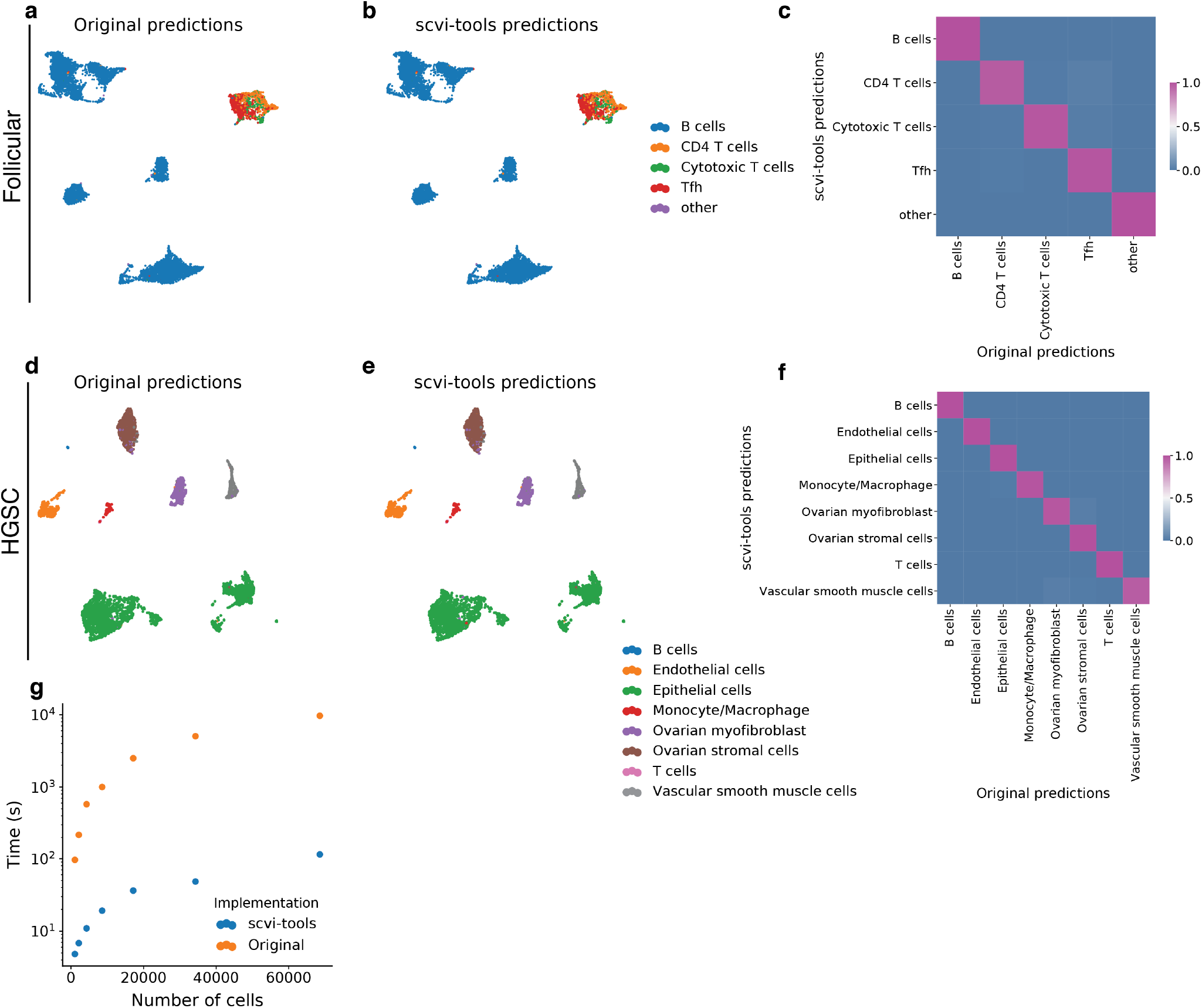
Evaluation of CellAssign implementation in scvi-tools. **a, b**, UMAP embedding of follicular lymphoma single-cell expression data, labeled by maximum probability assignments from the **(a)** original CellAssign implementation and **(b)** scvi-tools implementation. **c**, Row-normalized confusion matrix of scvi-tools predicted labels (rows) and study-derived cell annotations (columns) for follicular lymphoma expression data. **d, e**, UMAP embedding of HGSC single-cell expression data, labeled by maximum probability assignments from the **(d)** original CellAssign implementation and **(e)** scvi-tools implementation. **f**, Row-normalized confusion matrix of scvi-tools predicted labels (rows) and study-derived cell annotations (columns) for HGSC single-cell expression data. **g**, Runtime on downsampled versions of 68k peripheral blood mononuclear cells dataset for the original and scvi-tools implementations.

## Supplementary Tables

**Supplementary Table 1:** Overview of models and functionality currently implemented in scvi-tools.

| Modality | Model | Tasks |
| --- | --- | --- |
| scRNA-seq | scVI [5]<br>LDVAE [50]<br>scANVI [12]<br>AutoZI [91]<br>Solo [42]<br>CellAssign [39] | Dimensionality reduction, removal of unwanted variation, integration across replicates, donors, and technologies, differential expression, imputation, normalization of other cell- and sample-level confounding factors<br>scVI tasks with linear decoder<br>Automated annotation, all scVI tasks<br>Gene-wise model selection for zero-inflation<br>Doublet detection<br>Marker-based automated annotation |
| CITE-seq | totalVI [21] | Dimensionality reduction, removal of unwanted variation, integration across replicates, donors, and technologies, differential expression, protein imputation, imputation, normalization of other cell- and sample-level confounding factors |
| scATAC-seq | PeakVI [24] | Dimensionality reduction, removal of unwanted variation, differential accessibility, imputation, normalization of other cell- and sample-level confounding factors |
| Spatial Transcriptomics | gimVI [52]<br>Stereoscope [40]<br>DestVI [41] | Imputation of missing genes in spatial data using scRNA-seq reference<br>Deconvolution of spatial transcriptomics profiles<br>Multi-resolution deconvolution of spatial transcriptomics profiles |
| Multiple | scArches [38] | Transfer learning for reference-query integration applied on top of peakVI, scVI, scANVI, totalVI |

**Supplementary Table 2:** Gene sets for multiple covariates Drosophila analysis.

| Cell Sex Genes | Cell Cycle Genes | Marker Genes |
| --- | --- | --- |
| lncRNA:roX1 | PCNA | Argk |
| lncRNA:roX2 | dnk | Nrt |
| Sxl | RnrS | Ten-a |
| msl-2 | RnrL | Ten-m |
|  | Claspin | wb |
|  | Mcm5 | Act57B |
|  | Caf1-180 | drl |
|  | RPA2 | mid |
|  | HipHop | nemy |
|  | stg | lms |
|  | Mcm6 | CG11835 |
|  | dup | Gyg |
|  | WRNexo | ara |
|  | Mcm7 | tok |
|  | dpa | kirre |
|  | CG10336 | NK7.1 |
|  | Mcm3 | fj |
|  | Mcm2 | beat-IIIc |
|  | RpA-70 | CG33993 |
|  | Chrac-14 | dpr16 |
|  | CG13690 | CG15529 |
|  | RPA3 | CG9593 |
|  | asf1 | beat-IIb |
|  | DNApol-alpha73 | robo2 |
|  | CycE | Ama |
|  | DNApol-alpha50 | fz2 |
|  | Kmn1 | elB |
|  | Lam | noc |
|  | Nph | nkd |
|  | msd5 | fng |
|  | msd1 | vg |
|  | ctp |  |
|  | Set |  |
|  | scra |  |
|  | Chrac-16 |  |
|  | ncd |  |
|  | Ote |  |
|  | pzg |  |
|  | HDAC1 |  |
|  | nesd |  |
|  | tum |  |
|  | CG8173 |  |
|  | aurB |  |
|  | feo |  |
|  | pav |  |
|  | CG6767 |  |
|  | sip2 |  |
|  | Det |  |
|  | Cks30A |  |
|  | CycB |  |
|  | B52 |  |

## Supplementary Note 1

In this note, we describe the implementation of CellAssign in scvi-tools. Unlike other external methods in scvi-tools, there are slight differences in the scvi-tools implementation. First we describe the statistical model of CellAssign, highlighting differences between the scvi-tools implementation and the original implementation. Then we evaluate the scvi-tools implementation against the original on two datasets used in the original publication [39].

### Preliminaries

CellAssign takes as input a scRNA-seq gene expression matrix *X* with *N* cells and *G* genes along with a cell-type marker matrix *ρ* which is a binary matrix of *G* genes by *C* cell types denoting if a gene is a marker of a particular cell type. A size factor *s* for each cell may also be provided as input, otherwise it is computed empirically as the total unique molecular identifier count of a cell. Additionally, a design matrix *D* containing *p* observed covariates, such as day, donor, etc, is an optional input.

### Generative process

CellAssign uses a negative binomial mixture model to make cell-type predictions. The cell-type proportion is drawn from a Dirichlet distribution,

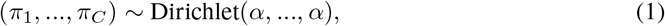

with *α* = 0.01.

CellAssign then posits that the observed gene expression counts *x*_*ng*_ for cell *n* and gene *g* are generated by the following process:

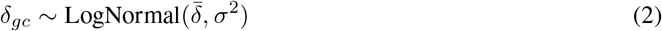

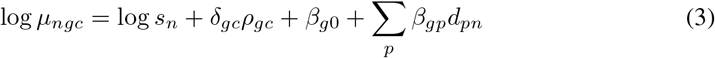

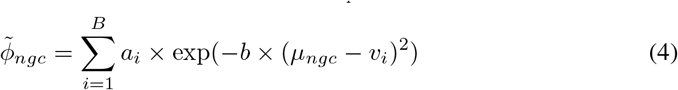

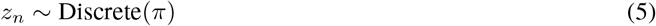

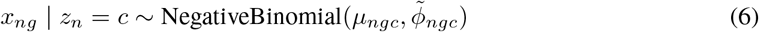

Notably, the logarithm of the mean of the negative binomial for cell *n*, gene *g*, given that it belongs to cell type *c* (Equation 3) is computed as the sum of (1) the base expression of a gene *g, β*_*g*0_, (2) a cell-type-specific overexpression term for a gene, *δ*_*gc*_, (3) an offset for the size factor, log *s*_*n*_, and (4) a linear combination of covariates in the design matrix (weighted by coefficients *β*_*gi*_). A cell-specific discrete latent variable, *z*_*n*_, represents the cell-type assignment for cell *n*.

Furthermore, the inverse dispersion of the negative binomial, 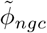 (Equation 4) is computed with a sum of radial basis functions of the mean centered on *v*_*i*_ with parameters *a*_*i*_ and *b*. In total, there are *B* centers *v*_1_, *v*_2_, …, *v*_10_ that are set a priori to be equally spaced between 0 and the maximum count value of *X*. Additionally, as in the original implementation we used *B* = 10. The parameters *a*_*i*_ and *b* are further described below.

### Inference

CellAssign uses expectation maximization, which optimizes its parameters and provides a cell-type prediction for each cell.

#### Initialization

CellAssign initializes parameters as follows: *δ*_*gc*_ is initialized with a LogNormal(0, 1) draw, 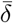 is initialized to 0, *σ*^2^ is initialized to 1, *π*_*i*_ is initialized to 1*/C, β*_*gi*_ is initialized with a 𝒩(0, 1) draw, and the inverse dispersion parameter log *a*_*i*_ is initialized to zero. Additionally, *b* is fixed a priori to be

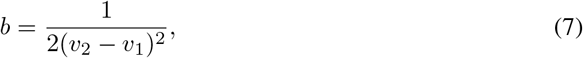

where *υ*_1_ and *υ*_2_ are the first and second center determined as stated previously.

#### E step

The E step consists of computing the expected joint log likelihood with respect to the conditional posterior, *p*(*z*_*n*_ | *x*_*n*_, *δ, β, a, π*), for each cell. This conditional posterior is computed in closed form as

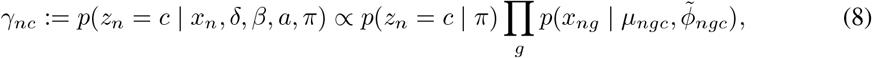

which is normalized over all *c*. The expected joint log likelihood over all *N* cells is then computed as

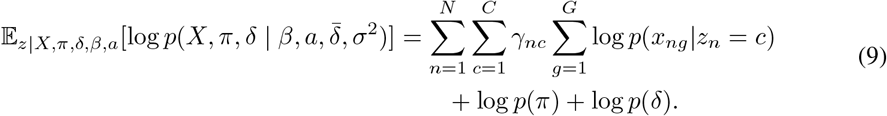

Herein lies the major difference between the scvi-tools implementation and the original CellAssign implementation. Notably, in scvi-tools we compute this expected joint log likelihood using a mini-batch of 1,024 cells, using the fact that

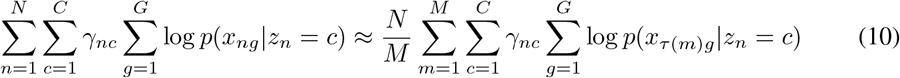

for a minibatch of *M* < *N* cells, where *τ* is a function describing a permutation of the data indices *{*1, 2, …, *N}*.

#### M step

The parameters of the expected joint log likelihood 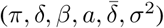 are optimized as in the original implementation, using the Adam optimizer [92], except that now an optimization step corresponds to data from one minibatch. Following the original implementation, we also clamped *δ >* 2. We also added early stopping with respect to the log likelihood of a held-out validation set.

### Evaluation

We downloaded two datasets with full metadata and cell-type marker matrices from the original publication [39] (from https://zenodo.org/record/3372746) and compared the scvi-tools implementation predictions to the original predictions for these datasets. The first dataset consisted of 9,156 cells from lymph node biopses of two follicular lymphoma (FL) patients. The second dataset consisted of 4,848 cells from a high-grade serous carcinoma (HGSC) patient. On both datasets, the scvi-tools implementation predictions were highly reproducible with the original implementation (Supplementary Figure 6a-f). UMAP embeddings used in Supplementary Figure 6 were the original embeddings from the publication and were retrieved from the downloaded objects.

Next, we evaluated the runtime of the two implementations on a desktop with an Intel Core i9-10900K 3.7 GHz processor, 2x CorsairVengeance LPX 64GB ram, and an NVIDIA RTX 3090 GPU. To do so, we used a dataset of 68k peripheral blood mononuclear cells from 10x Genomics [78], and used the same cell-type marker matrix as the FL dataset (23 markers, 5 cell types). Across a range of cells, we observed that the scvi-tools implementation is consistently faster and thus more scalable (Supplementary Figure 6g). This is mostly due in part to the minibatching, which is a capability present in the original codebase, but not set as default (also no guidance on how to set it). Thus, we have shown that the scvi-tools implementation of CellAssign is both reproducible, and by default, more scalable than the original implementation.

